# Versatile cell-based assay for measuring DNA alkylation damage and its repair

**DOI:** 10.1101/2021.02.25.432902

**Authors:** Yong Li, Peng Mao, Evelina Y. Basenko, Zachary Lewis, Michael J. Smerdon, Wioletta Czaja

## Abstract

DNA alkylation damage induced by environmental carcinogens, chemotherapy drugs, or endogenous metabolites plays a central role in mutagenesis, carcinogenesis, and cancer therapy. Base excision repair (BER) is a conserved, front line DNA repair pathway that removes alkylation damage from DNA. The capacity of BER to repair DNA alkylation varies markedly between different cell types and tissues, which correlates with cancer risk and cellular responses to alkylation chemotherapy. The ability to measure cellular rates of alkylation damage repair by the BER pathway is critically important for better understanding of the fundamental processes involved in carcinogenesis, and also to advance development of new therapeutic strategies. Methods for assessing the rates of alkylation damage and repair, especially in human cells, are limited, prone to significant variability due to the unstable nature of some of the alkyl adducts, and often rely on indirect measurements of BER activity. Here, we report a highly reproducible and quantitative, cell-based assay, named alk-BER (alkylation Base Excision Repair) for measuring rates of BER following alkylation DNA damage. The alk-BER assay involves specific detection of methyl DNA adducts (7-methyl guanine and 3-methyl adenine) directly in genomic DNA. The assay has been developed and adapted to measure the activity of BER in fungal model systems and human cell lines. Considering the specificity and conserved nature of BER enzymes, the assay can be adapted to virtually any type of cultured cells. Alk-BER offers a cost efficient and reliable method that can effectively complement existing approaches to advance integrative research on mechanisms of alkylation DNA damage and repair.

---

DNA alkylation induced by methylating agents such as environmental carcinogens (e.g., smoke), by-products of cellular metabolism (e.g., methyl group donor S-adenosyl methionine), or chemotherapy drugs (e.g., temozolomide, procarbazine) represents one of the most abundant types of DNA base damage that forms in human cells. Monofunctional alkylating agents, like methyl methanesulfonate (MMS) or the anticancer drug temozolomide (TMZ) induce formation of N-methyl and O-methyl DNA adducts such as N7-methylguanine (7meG), N3-methyladenine (3meA) and O6-methylguanine (O6meG)^1,2^. Methyl DNA adducts (MDAs) have cytotoxic and mutagenic properties because of their ability to block gene transcription and interfere with the fidelity of DNA replication. Persistent and inefficiently repaired methyl DNA adducts can induce microsatellite instability, frameshift mutations, and G → A transition mutations, that are commonly found in genes critical for malignant transformation, including the *H-ras* oncogene or *TP53* tumor suppressor gene^3–5^. Despite their carcinogenic properties, DNA alkylating agents, such as dacarbazine, temozolomide and streptozotin, have been used for decades in treating various cancers, including melanoma, glioma, and lymphoma^2,6,7^. Therapy with alkylating agents can be effective; however, these agents are extremely toxic and prolonged treatment often leads to chemoresistance and formation of secondary cancers^8,9^. Human responses to alkylating agents vary considerably between individuals, which highlights the involvement of genetic and epigenetic mechanisms in the modulation of cellular toxicity to alkylating agents^2,10,11^.

Base excision repair (BER) is the primary pathway involved in the removal of alkylation DNA damage induced by methylating agents^2,12,13^. Repair of methyl DNA adducts through the BER pathway is accomplished in four sequential steps, each carried out by a specific group of enzymes^14^. The first step is catalyzed by DNA glycosylases (e.g., AAG), which specifically recognize and bind to a damaged base, and subsequently catalyze cleavage of the glycosidic bond between the damaged base and DNA backbone^15,16^. This reaction results in the release of the damaged base from DNA, and the formation of abasic AP (apurinic/apyridiminic) sites. The second step involves incision of the DNA backbone 5’ upstream of the AP sites by AP endonucleases (e.g., APE1), which results in the formation of single strand DNA breaks (SSBs)^17,18^. In mammalian cells, AP sites and SSBs are recognized by poly ADP-ribose polymerase 1 (PARP1)^19,20^. Activated PARP1 catalyzes the formation of ADP-ribose chains, which serve as a docking platform that facilitates recruitment and assembly of the multiprotein BER complex (XRCC1-POLβ-LIG3). Breaks in DNA are filled in by DNA polymerases (primarily POLβ) using the undamaged complementary DNA strand as a template^21^. Nicks in the damaged strand are sealed by ligases (e.g., LIG3), which finalizes repair of the damaged DNA strand^22,23^.

Importantly, methylation-derived repair intermediates such as AP sites and SSBs are highly cytotoxic and mutagenic. Therefore, individual steps in the BER pathway need to be tightly regulated and coordinated to prevent accumulation of those intermediates, cell death, mutagenesis, and carcinogenesis^2,24^. Genetic studies in yeast, mouse models, and human cells have demonstrated that loss of the tight coordination between individual steps in the BER pathway can trigger genome instability, increased mutagenesis, or cell death^2,5,24–27^. Levels and activities of BER proteins vary significantly between cells, tissues, and individuals and correlate with cancer risk and response to alkylation chemotherapy^2,10,11,25,28–32^. Therefore, measuring and understanding differences in the rate of BER upon alkylation DNA damage could contribute to the development of new approaches in personalized disease prevention and treatment.

The BER pathway is dysregulated in many cancers and is often associated with cancer heterogeneity, metastasis, and chemoresistance. Pharmacological inhibition of BER with PARP inhibitors (e.g., olaparib) has shown enhanced cytotoxicity of various anticancer agents, especially in tumors with defects in homologous recombination^33–37^. Identifying a pre-existing BER imbalance within a tumor may be highly relevant for determining whether therapy involving PARP inhibitors and alkylating agents can be beneficial.

Quantitation of DNA adduct formation and repair has greatly advanced our understanding of DNA repair processes. A number of methods have been developed for quantitative analysis of various enzymatic steps and the overall capacity of BER to repair alkylation DNA damage. The most sensitive methods for detection and quantitation of alkyl DNA adducts include HPLC/^32^P-postlabeling, mass spectrometry-based adductomics, and radiolabeling^38–41^. These methods offer high sensitivity; however, they require specialized equipment, expertise, and complex sample preparation, which hinders the convenient use of those approaches to investigate cellular BER mechanisms.

The most commonly used cell-based methods to investigate BER, include comet assays and host cell reactivation (HCR) assays. The comet assay is a single-cell electrophoresis technique that can be used to assess the capacity of BER to repair alkylation DNA damage when performed under alkaline conditions^42^. This assay can be used to analyze total levels of BER repair intermediates, such as alkali labile sites (e.g., abasic sites) and single strand breaks; however, it does not quantitatively distinguish between these intermediates. The standard comet assay may not reliably detect persistent and intact base modifications (e.g., 7meG or 3meA) that are not converted to AP sites or SSBs. In addition, the comet assay may not detect lesions that form and persist within highly inaccessible heterochromatin fractions of the genome. Furthermore, the standard comet assay workflow is laborious and prone to day-to-day variability. It may also require extensive optimization of experimental conditions, including pH or salts used during the alkaline electrophoresis steps, to achieve sensitivity and consistent reproducibility^42,43^. HCR is another method that has been used to measure the capacity of BER to repair alkylation DNA damage in living cells^44,45^. HCR relies on the transfection of the non-replicating DNA plasmid with a reporter gene (e.g., *luciferase*) that contains chemically-induced DNA base damage, which is subject to repair by the BER pathway. The presence of the DNA base damage within the reporter gene inhibits its expression, whereas the repair of base damage re-activates reporter expression. The HCR assay can be especially challenging to assess repair of alkylation DNA damage, due to in vitro instability of the alkyl DNA adducts (e.g., 7meG and 3meA), which can markedly affect assay reproducibility^45^. Also, HCR involves a non-genomic DNA substrate that does not necessarily reflect the complexity of the genomic chromatin environment.

More recently, high resolution, high throughput approaches such as LAF-seq (Lesion-Adjoining Fragment Sequencing) or NMP-seq (N-methyl purine sequencing) utilizing next generation DNA sequencing have been developed to enable precise mapping and quantitation of methyl DNA adducts across the genome, and at specific genomic loci^46,47^. These approaches offer unprecedented single base resolution, but can be laborious, involving generation of DNA sequencing libraries and extensive bioinformatics analyses of the sequencing data, especially when used with human cells. In addition, those methods may require high (non-physiological) doses of DNA damaging agents and large amounts of input DNA.

Here we report a reliable, gel-based method, called alk-BER, that offers a fast and quantitative measure of BER capacity in living cells. Alk-BER was developed by an adaptation of a previous methods for UV-induced DNA damage quantitation by alkaline gel electrophoresis, originally developed by Sutherland et al.^48^. We developed a set of optimized protocols to analyze formation and repair of alkylation-induced DNA damage in various model organisms, including fungal cells (*S. cerevisiae, N. crassa*) and human cancer cell lines.

Alk-BER can be used to assess overall capacity of BER to repair MMS-induced alkylation DNA damage within the genome of living cells. It should be noted that alk-BER assay does not directly measure activity of the endogenous BER enzymes in the repair of DNA alkylation damage, but it measures changes in the levels of DNA alkylation in the genomic DNA that strongly correlate with the capacity of the BER pathway to directly and specifically repair DNA alkylation in the genomic DNA. The assay can be used to facilitate identification of new, conserved regulators of the BER pathway by using complementary model eukaryotic systems, including fungal model organisms and human cells. Application of the alk-BER assay could also facilitate identification of BER-deficient cancer sub-types, which might represent suitable targets for therapy with alkylating agents and/ or PARP inhibitors.

## Materials and methods

### DNA damage and time course of repair in yeast cells

Yeast (*S. cerevisiae*) liquid cell cultures were inoculated from single colonies and grown in 10 ml of YPD (Yeast extract -Peptone-Dextrose) medium for ∼ 16 h at 30 °C in an orbital shaker. The following day, cells were sub-cultured in fresh media and grown until the cultures reached the logarithmic stage of growth, as determined by measuring the optical density of the cell culture (e.g., OD_600_ ∼ 0.6). Next, MMS was added to the liquid cultures at a final concentration of 20 mM and cells were incubated for 10 min at 30 °C in an orbital shaker. Alternatively, cells were treated with 3.5 mM MMS for 1–3 h, followed by removal of media containing MMS and repair time course in fresh media for 1–6 h. Cells were then harvested by centrifugation, and the supernatant fractions containing MMS were removed and disposed following DEHS guidelines. Cell pellets were washed with ice-cold phosphate buffered saline (1X PBS), re-suspended in a pre-warmed YPD media, and allowed to repair for a total of 3 h. Extended repair time points (longer than 4–5 h) were avoided to ensure that restoration of the genome integrity was due to the activity of BER, and not due to lesion bypass and DNA replication. Cultures were incubated with continuous shaking and cell aliquots were collected at different repair time points^49,50^.

### Yeast genomic DNA isolation

Yeast genomic DNA was extracted with the glass bead method following previously established protocols^51^. Briefly, cell pellets were mixed with 250 μL of DNA lysis buffer [2% (vol/vol) Triton X-100, 1% SDS, 100 mM NaCl, 10 mM Tris·HCl, pH 8.0, 1 mM EDTA, 250 μL of PCI (phenol: chloroform: isoamyl alcohol = 25:24:1), and 150 μL of acid-washed glass beads, and vortexed vigorously for 4 min. Next, 200 μL of 1XTE buffer (10 mM Tris·HCl, pH 7.5, 1 mM EDTA) were added and cell lysates were centrifuged at 14,000 rpm at 4 °C. The supernatant fraction was transferred to a fresh Eppendorf tube and mixed with 1 mL of ice-cold 200 proof ethanol. Samples were incubated at − 80 °C for 15 min to facilitate formation of the DNA precipitate. Next, samples were centrifuged at 14,000 rpm at 4 °C, and washed with 70% (vol/vol) ethanol. The DNA pellets were dissolved in 200 μL of 1XTE buffer and incubated with 2 μL of RNase A (Thermo Fisher Scientific, cat # EN0531) at 37 °C for 1 h to remove RNA. DNA was subsequently ethanol-precipitated, dissolved in sterile deionized H_2_O, and then stored at − 80 °C.

### AAG and APE1 reactions

Purified genomic DNA was processed with (+) or without (−) an enzymatic cocktail composed of AAG (New England BioLabs, cat# M0313S) and APE1 (New England BioLabs, cat# M0282), to convert MMS-induced 7meG and 3meA to SSBs. DNA samples (0.6–1 μg) were incubated with 1 μL of AAG and 1 μL of APE1 in the reaction buffer (70 mM MOPS, pH 7.5,1 mM dithiothreitol (DTT), 1 mM EDTA, 5% glycerol) at 37 °C for 1 h in a total reaction volume of 20 μL^52^. Methylated bases were cleaved by AAG glycosylase and the resulting abasic sites were cleaved by APE1 endonuclease. The reactions were stopped by adding DNA loading buffer (50 mM NaOH, 1 mM EDTA, 3% Ficoll, 0.025% Bromocresol green, 0.041% Xylene cyanol). Following enzymatic digestion of DNA cleavage products, single and double strand breaks were resolved on 1.2% alkaline agarose gels and stained with SYBR Gold^49,50^.

### Alkaline agarose gel electrophoresis

Alkaline agarose gel electrophoresis was performed following previously published protocols with modifications^48,53^. The large gel electrophoresis box and casting tray were used. The agarose solution was prepared by adding 4.3 g of agarose to 360 ml of H_2_0 in a 1 L Erlenmeyer flask and then heating in a microwave oven until the agarose was dissolved. The solution was cooled to 55 °C and then followed by addition of a 0.1 volume (40 ml) of 10X alkaline agarose gel electrophoresis buffer (500 mM NaOH, 10 mM EDTA, pH 8.0). Addition of 10X alkaline buffer to a hot agarose solution should be avoided because NaOH in the buffer may cause hydrolysis of the agar. The agarose solution was poured into a large gel casting tray. After the gel was completely solidified, it was mounted in the electrophoresis tank. Then the tank was filled with 1X alkaline electrophoresis buffer until the gel was covered with the buffer at a depth of 3–5 mm above the gel. DNA samples after AAG and APE1 digestion were collected and 6X alkaline gel-loading buffer (300 mM NaOH, 6 mM EDTA, 18% Ficoll, 0.15% Bromocresol green, 0.25% Xylene cyanol) was added to each sample. Chelating all Mg^2+^ with EDTA (component of the 6X alkaline gel-loading buffer) is important before loading the samples onto the alkaline agarose gel because in solutions with a high pH, Mg^2+^ can form insoluble Mg (OH)_2_ precipitates that entrap DNA and inhibit DNA mobility through the gel. Samples were loaded and the gel was run at room temperature at 30 V for 19–24 h. Alternatively, the electrophoresis can be run at 4 °C in a cold room. We found that running the gel in a cold room helped improve the sharpness of the DNA bands. Note that after the run is completed, the Bromocresol green dye may not be visible because of dye diffusion in the gel. The gel was transferred to a large plexiglas tray, covered with neutralizing solution (1 M Tris–HCl pH 7.6, 1.5 M NaCl), and incubated for 30 min with gentle shaking on an orbital shaker. Next the gel was transferred to staining solution (1X TAE buffer with SYBR gold) and stained for 1 h with gentle shaking. The container with the gel was covered with aluminum foil to protect the staining solution from light. Following staining, the gel was rinsed and de-stained with H_2_0 for 30 min with gentle shaking.

### Equipment and settings

Images of the gels were acquired using the AI600 Imager (Amersham), by performing fluorescence Epi-RGB blue light (460 nm), filter Cy2 capture with automatic exposure.

### Quantitative analysis of DNA damage and repair following MMS treatment

Quantitation of methylated bases in genomic DNA was performed on phosphor image data by using ImageQuant TL 8.2 software (Cytiva) and number-average DNA length analysis. The number-average length of genomic DNA (±AAG/ APE treatment) was used to calculate the average number of SSBs/kb, and the percentage of MDAs removed (% repair) was determined as described previously^49,51^. Briefly, using the ImageQuant software functions, each data point on the gel image is marked with a box encompassing the entire length of the lane to give the total area of each lane. The data point corresponding to 1/2 the total area, designated as *X*_*med*_, is then determined. The *X*_*med*_ value indicates the median migration distance of the DNA fragments. *X*_*med*_ is converted to the median length *L*_*med*_ of DNA molecules by using a standard curve generated from the migration of DNA-size markers. The number average molecular length, *L*_*n*_, is calculated from *L*_*med*_ by using the equation *L*_*n*_ = 0.6 *L*_*med*_^54^, assuming a Poisson distribution of DNA fragments. Numbers of SSBs/kb is calculated using the following equation; SSBs/kb = 1/L_*n*_ (+ AAG + APE1) − 1/L_*n*_ (-AAG-APE1). Calculated numbers of SSBs per kb DNA fragment indicate numbers of MDAs.

### DNA damage and time course of repair in multicellular fungus *Neurospora crassa*

Liquid cultures of *N. crassa* were grown for 11 h in VMM (Vogel’s minimal medium) media in an orbital shaker at 30 °C. MMS was added to liquid cultures at a final concentration of 3.5 mM. Cells were incubated in the presence of MMS for 3 h. Next, the cells were collected by using a Buchnner funnel, and washed with 500 ml of VMM media to remove MMS. Washed cells were transferred to pre-warmed VMM media and then allowed to grow at 30 °C for 4 h to enable repair of damaged DNA. Aliquots of cells were harvested and immediately frozen in liquid nitrogen. Genomic DNA was isolated and 300 ng of genomic DNA was digested with APE1 endonuclease (cat # M0282S, New England Biolabs), AAG glycosylase (cat # M0313S, New England Biolabs), or both enzymes in MOPS reaction buffer (70 mM MOPS, pH 7.5,1 mM dithiothreitol DTT, 1 mM EDTA, 5% glycerol) for 1 h and 15 min at 37 °C. Reactions were stopped by adding alkaline DNA loading buffer (50 mM NaOH, 1 mM EDTA, 3% Ficoll). Samples were resolved on a 1.2% alkaline agarose gel. Agarose gels were run in a cold room at 25 V for 17 h and then incubated in neutralization buffer (1.5 M NaCl, 1 M Tris–Cl pH 7.6) for 45 min before being stained with SYBR Gold (cat # S-11494, Life Technologies) for 40 min and de-stained for 30 min before imaging^55^.

### DNA damage and time course of repair in human cells

Human adrenal carcinoma cells (SW13) were a gift from Dr. Feng Gong laboratory at the University of Miami. Human fibroblast CHON-002 were a gift from Dr. Rafael Contreras laboratory at the Hormel Institute, University of Minnesota. Human haploid HAP1 cells were purchased from Horizon Discovery Biosciences. SW13 cells were cultured in DMEM (Sigma, cat# D5796) supplemented with 10% fetal bovine serum FBS (Gibco, cat# 26140-079). Human fibroblast CHON-002, leukemia HAP1, and lymphoblastoid cells GM12878 were cultured according the ATCC guidelines. Cell cultures were routinely tested for mycoplasma by using a mycoplasma detection kit (ATCC, cat# 30-1012 K). Cell doubling time was determined following ATCC guidelines and viability was routinely monitored with trypan blue. Cells were seeded in T25 flasks (∼ 500,000 cells per dish) and grown for 16–24 h until cells reached 60–70% confluence. One T25 flask was set for each data point to be collected. In DNA damage dose response experiments SW13 cells were treated with (0-20 mM) MMS in 1XPBS for 10 min at RT (room temperature). In DNA damage and repair experiments, SW13 cells were treated with 10 mM (0.1%) MMS for 10 min in 1xPBS at RT, or alternatively ice-cold, serum-free media can be also used for MMS treatments. Ice-cold treatment is used to inhibit endogenous background BER during the MMS treatment. MMS was removed and cells were washed with 1X PBS. Next, fresh pre-warmed media were added and cells were allowed to repair for 0, 3, 8, or 22 h. GM 12878 cells were treated with 5 mM (0.05%) MMS for 5 min. Total genomic DNA was purified from each time point using the PureLink genomic DNA mini kit (K 182001, Thermo Fisher Scientific). DNA was processed with AAG and APE1 enzymes as described above. DNA was resolved on a 1.2% alkaline agarose gel, run at 30 V for 22 h, stained with SYBR gold, and quantified as described above.

### Western Blotting

HAP1, SW13 and CHON-002 cells were harvested and frozen at − 80 °C. Protein extracts were prepared in a RIPA buffer (Santa Cruz, sc-24948) plus phosphatase inhibitor (Santa Cruz, sc-45044 and sc-45044) and equal amounts of protein were separated on TGX stain-free (Bio-Rad, cat#5678083). Proteins were transferred onto TransBlot LF PVDF (Bio-Rad) and analyzed by Western blotting using antibodies recognizing the following proteins: AAG (Abcam ab155092, 42KD, 1:5000), beta-actin (Sigma A5441, 33KD, 1:5000).

## Results

### Alk-BER assay workflow

The conserved nature of the BER pathway enables easy adaptation of the assay to various cell types and model organisms from lower eukaryotes to human cells. We initially developed the alk-BER assay to measure BER in yeast cells (*S. cerevisiae*) and successfully adapted the assay to other fungal model organism (*N. crassa*) and human cells. The alk-BER assay is based on the enzymatic conversion of the MMS-induced damaged DNA bases to single strand breaks (SSBs), which are subsequently resolved on alkaline agarose gels and quantified. The rate of BER can be analyzed by performing a DNA damage and time course for repair and assessing the rate of removal of damaged bases from the total genomic DNA. The assay is performed in 5 simple steps (Fig. 1) and can be completed in 3 days. The first step involves exposing the cells to sub-lethal doses of MMS to induce formation of methyl DNA adducts (mainly 7meG and 3meA). In the second step, cell aliquots corresponding with DNA damage and repair time points are collected and total genomic DNA is isolated. The third step involves conversion of MDAs to SSBs with BER enzymes (AAG glycosylase and APE1 endonuclease) that specifically bind and cleave the DNA at sites of MDAs. The fourth step involves running the samples on alkaline agarose gels to separate DNA fragments containing SSBs from the bulk genomic DNA that does not contain damage. The final step involves staining of the separated DNA fragments in the gel, acquiring an image of the gel and performing quantitation of MDAs. The alk-BER method directly quantifies the numbers of MDAs in purified genomic DNA and analyzes the kinetics of DNA repair at the whole-genome level. In order to assess the capacity of BER using the alk-BER assay, cells should be exposed to sub-lethal doses of MMS, that induce detectable levels of MDAs, but do not induce substantial cell death. To determine sub-lethal doses for specific cell line it is recommended to perform MMS dose response experiment encompassing range of MMS doses from low to high. Cell death is induced when unrepaired lesions persist due to inability of BER to efficiently repair abnormally high levels of DNA damage or when BER activity is compromised (e.g., BER gene mutants). BER capacity should also be analyzed within a specific, experimentally determined window of time following DNA damage, typically 0–3 h for yeast cells, or 0–24 h for human cells, to avoid interference from lesion bypass and DNA replication.

**Figure 1.**
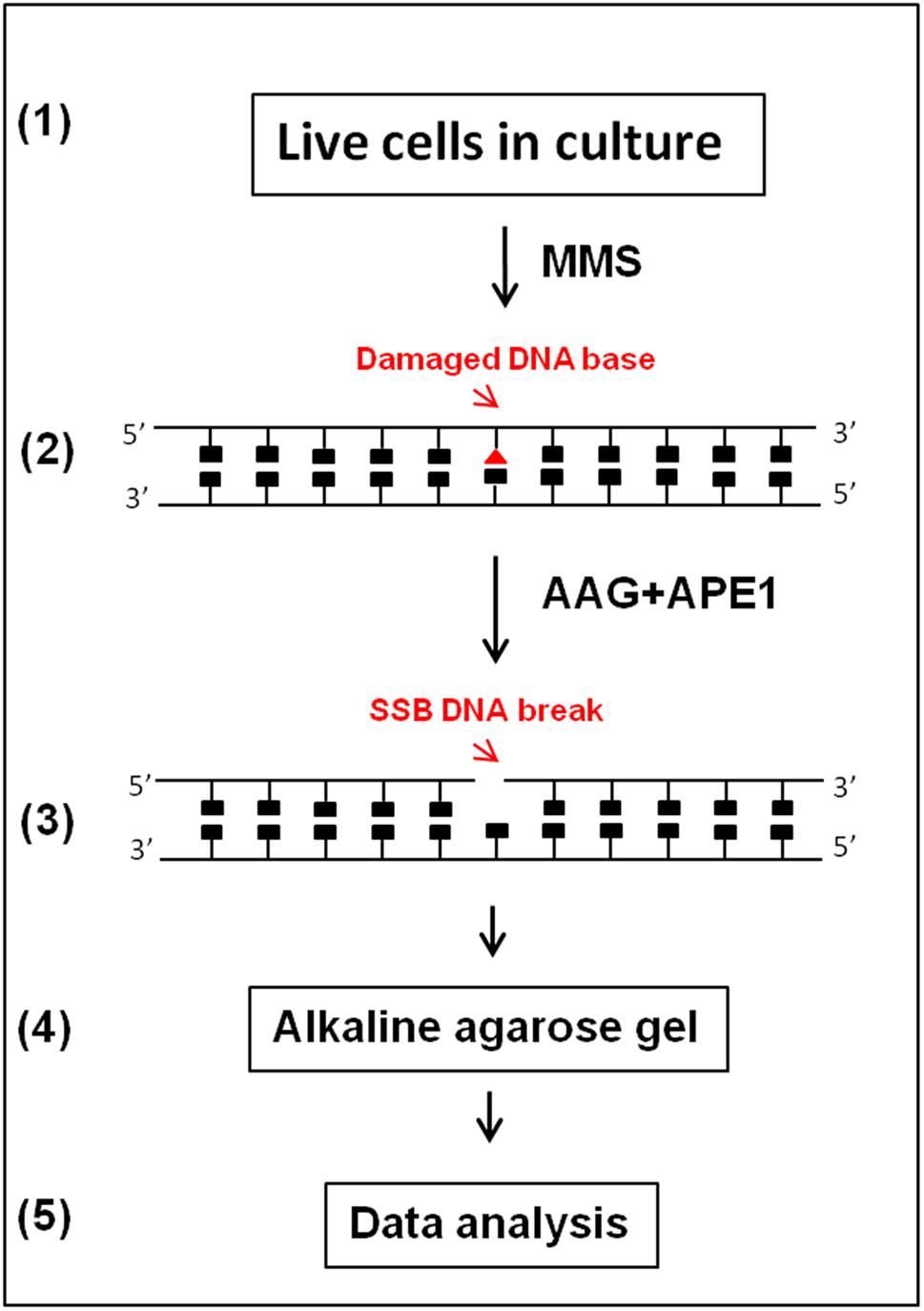
Schematic outline of alk-BER assay. The assay involves exposing the cells to MMS (step 1), isolation of total genomic DNA (step 2), conversion of MMS-induced methylated bases to SSBs with damage specific enzymes AAG and APE1 (step 3), separation of DNA fragments containing SSBs by alkaline agarose gel electrophoresis (step 4), gel staining, imaging, and quantitation of MDAs (step 5).

### Alk-BER in fungal cells

A DNA damage dose response assay was performed by exposing yeast cells (strain BY4741) to increasing doses of MMS (5, 10, 20 mM) for 10 min at 30 °C. Total genomic DNA was isolated and treated with AAG and APE1 enzymes. DNA samples were resolved by alkaline agarose gel electrophoresis and numbers of MDAs were quantified as described above. As the gel is run at alkaline pH, hydrogen bonding between the two DNA strands is broken to facilitate separation of strands containing breaks from non-damaged genomic DNA. The denatured DNA is maintained in a single-stranded state and migrates through the alkaline gel as a function of its size, forming a distinct smear. Increased formation of MMS-induced DNA lesions in response to increased doses of MMS is shown as increases in lower molecular weight DNA molecules, also visible as a smear (Fig. 2A). The frequency of MDAs (7meG and 3meA) was calculated and plotted as the number of methyl A, G per kb fragment as a function of increasing doses of MMS (Fig. 2B). The proportional relationship between increasing MMS doses and numbers of MDAs indicates a high sensitivity of the alk-BER method, ∼ 1.0 MDAs per 10,000 bases, that is induced by 5 mM MMS dose, and ∼ 4.0 MDAs per 10,000 bases induced by 20 mM MMS in yeast cells (Fig. 2B).

**Figure 2.**
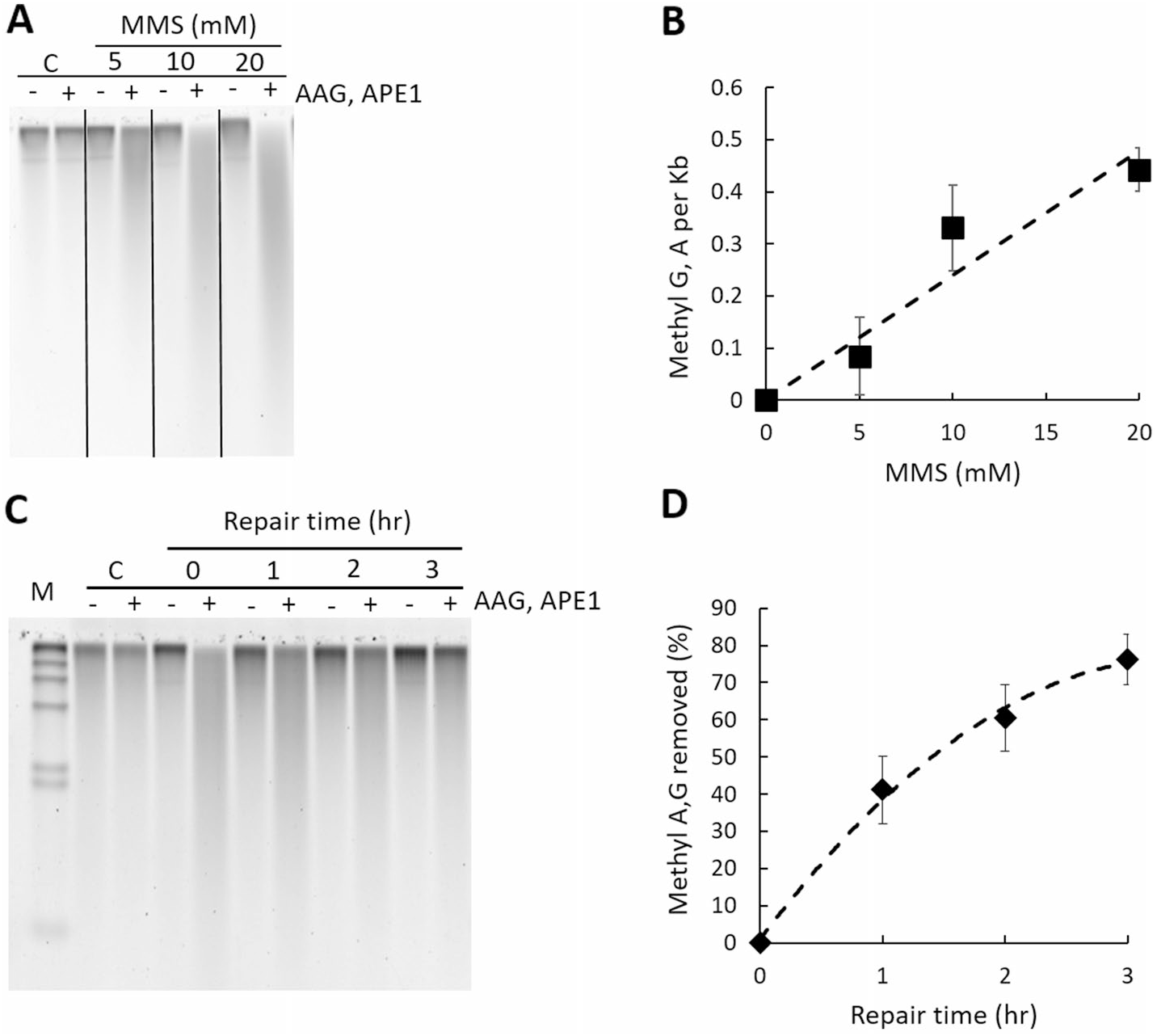
Alk-BER assay in yeast cells (*S. cerevisiae*). (**A**) Representative alkaline agarose gel image of MMS-induced DNA damage dose response in the BY4741 strain of *S. cerevisiae*. Genomic DNA of cells not exposed to MMS (C: control), and DNA of cells exposed to increasing doses of MMS (5, 10, or 20 mM) was resolved on alkaline agarose gel. Each DNA sample was treated with (+) and without (−) a cocktail of AAG and APE1 enzymes. (**B**) Dose dependent increase in the numbers of MMS-induced methyl G, A per 1 kb DNA fragment. Each data point denotes the average value and standard deviation of three independent experiments. (**C**) Representative gel image of DNA damage and repair time course in the BY4741 strain of *S. cerevisiae*. M: DNA size standard lambda/HindIII. C: control, cells not exposed to MMS, 0: cells collected after 10 min exposure to 20 mM MMS, 1–3 h: cells collected after 1, 2, 3 h of repair. (**D**) Quantitative representation of data displayed in panel C. Formation and repair of MMS-induced methyl G and A (7meG, 3meA), as a function of repair time. Each data point represents an average of 3 independent experiments; error bars were calculated based on standard deviation. Gel image presented in panel (**A**) has been cropped. Original, uncropped gel image is included in the supplementary data.

DNA damage and time course of repair has been performed to evaluate the overall rate of BER to repair alkylation DNA damage in the yeast wild type strain BY4741. DNA molecules containing MMS-induced MDAs are converted to SSBs. Proficient DNA repair and restoration of genome integrity can be visually monitored as progressive shortening of the DNA smear in migration and restoration of the genome integrity by formation of high molecular weight DNA. DNA was processed with AAG and APE1 and resolved on alkaline agarose gels as described previously. A representative gel image demonstrating DNA damage and repair in the BY4741 yeast strain is shown (Fig. 2C) and corresponding quantitative analysis of the gel is also shown (Fig. 2D). BER in the WT BY4741 yeast strain is proficient, and over 80% of the total MDAs in the genome are repaired after 3 h repair time at the dose used (Fig. 2D). The specificity of the alk-BER method was validated using yeast and *Neurospora* mutant cells deficient in BER. The yeast *mag1Δ* mutant has no Mag1 glycosylase (orthologue of human AAG) and is deficient in cleaving MMS-induced 7meG and 3meA from the DNA. Cells from the WT and *mag1Δ* strains were exposed to MMS for a total of 3 h followed by MMS removal and a 6 h-long repair time course to allow cells to repair damaged DNA. During the 3 h-long MMS exposure, mutant cells displayed higher levels of MDA formation as compared to WT because endogenous BER is inactive in the mutant cells, which results in accumulation of MDAs under the conditions of continuous MMS exposure. After removal of MMS, WT cells with proficient BER were able to clear most of the lesions during the repair time course, whereas BER-deficient *mag1Δ* cells contained high levels of unrepaired MDAs (Fig. 3A–C). The alk-BER assay was successfully adapted to multicellular fungal model *Neurospora crassa*. DNA damage and repair time course was performed with *Neurospora* WT and *mag1Δ* cells^55^. *Neurospora* cells of the wild type laboratory strain were exposed to 3.5 mM MMS continuously for 3 h, followed by repair time course in media without MMS for 5 h. Genomic DNA was isolated and processed with combinations of AAG and APE1 enzymes and resolved on alkaline agarose gel. BER capacity to repair MMS-induced lesions in *Neurospora* cells is very efficient, with nearly complete restoration of genome integrity following 2 h repair period (Fig.S2).

**Figure 3.**
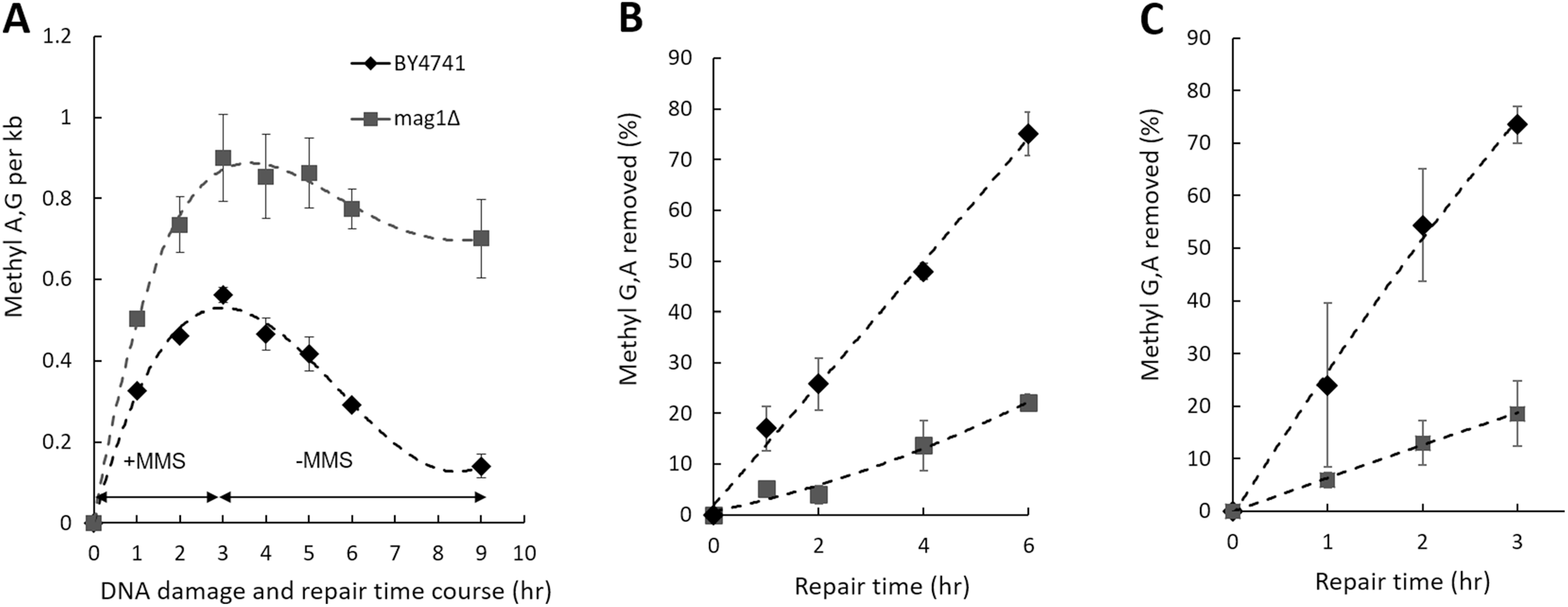
Alk-BER assay validation with BER-deficient yeast mutant cells, *mag1Δ*. (**A**) The BER rate was analyzed in *mag1Δ* mutant. WT and mutant cells were treated with 3.5 mM MMS for 1–3 h, followed by removal of media containing MMS and repair time course in fresh media for 1–6 h. (**B**) Repair rates expressed as a function of % of methyl A, G removed over the repair time. (**C**) Cells were treated with 20 mM MMS for 10 min followed by repair time course for total of 3 h. Repair rates expressed as a function of % of methyl A, G removed over the repair time. Each data point represents an average of two independent experiments.

### Alk-BER in human cells

The human adrenal carcinoma SW13 cell line was used to adapt and optimize the alk-BER assay for assessing rates of BER in human cells. A series of MMS-dose response experiments were initially performed to determine the appropriate range of MMS concentration and time of the exposure for induction of detectable levels of MDAs at sub-lethal MMS doses. Cell viability was routinely monitored with trypan blue^56^. Representative MMS dose response data are presented in Fig. 4A,B. Increases in the smear length in response to increasing doses of MMS in the (+ AAG&APE1) lanes indicate enhanced formation of MDAs. The minimal smear in (− AAG&APE1) lanes reveals formation of MDA-derived BER intermediate AP sites and SSBs that form in DNA as a result of the continuous activity of endogenous BER during MMS exposure. AP sites are fragile in alkaline conditions and can spontaneously convert to SSBs contributing to the smear in (-enzyme) lanes. The proportional relationship between increasing MMS doses and numbers of MDAs indicates a high sensitivity of the alk-BER method, ∼ 0.7 MDAs per 10,000 bases with 5 mM MMS dose in human cells. Efficiency of the individual enzymes, AAG and APE1 to convert MMS-induced MDAs to SSB in genomic DNA was assess. We found that the cocktail of both enzymes AAG and APE1 works most efficient in converting MDAs to SSBs (Fig. 4C). The BER capacity in SW13 cells was analyzed by performing DNA damage and a time course of repair as described in the methods section. Cells were exposed to 10 mM MMS for 10 min in 1X PBS at RT, followed by removal of MMS and repair for 22 h in fresh media. The MMS dose used was a sublethal dose, that did not trigger significant cell death, as demonstrated by the cell viability data (Fig. 4F). After 22 h post MMS exposure nearly 70% of the genome was restored in SW13 cells. Representative data showing the image of the alkaline agarose gel and data quantitation are presented (Fig. 4D,E). Other human cell lines were subject to alk-BER assay, including untransformed fibroblast cells CHON-002, and leukemia cancer cells HAP1. The BER capacity to remove MMS-induced adducts was quantitated over distinct repair time points. The removal of MDAs appears very slow (∼ 0–30% repair) during the first 0-8 h post MMS exposure and is consistently observed in many different human cell lines we tested. Interestingly, the repair rates vary significantly between different cell lines, and unlike in fungal cells, the rates do not appear to correlate well with the levels of AAG enzyme in the panel of cell lines we tested (Fig. 5A,B). Additionally, alk-BER assay was performed with human lymphoblastoid cell line GM12878. Cells were exposed to 5 mM MMS for 5 min, and repair rate was analyzed at 2.5 and 5 h in the repair time in media without MMS. Clearly, very slow repair was detected during the initial 5 h repair (Fig. S3). Specificity of alk-BER was analyzed by exposing SW13 cells to increasing doses of oxidative agent hydrogen peroxide (H2O2), and temozolomide (TMZ), clinically used SN1 alkylator, known to induce 7meG, 3meA and O6meG adducts^57^. Exposure to H2O2 does not result in formation of a dose-dependent DNA smear, indicating that alk-BER assay is specific to methyl adducts, and it cannot detect oxidative DNA damage (Fig. S1A). As expected, alk-BER can specifically detect TMZ-induced DNA methyl adducts in a dose dependent manner (Fig. S1B,C), although the TMZ potency to induce MDAs appears lower than the potency of MMS.

**Figure 4.**
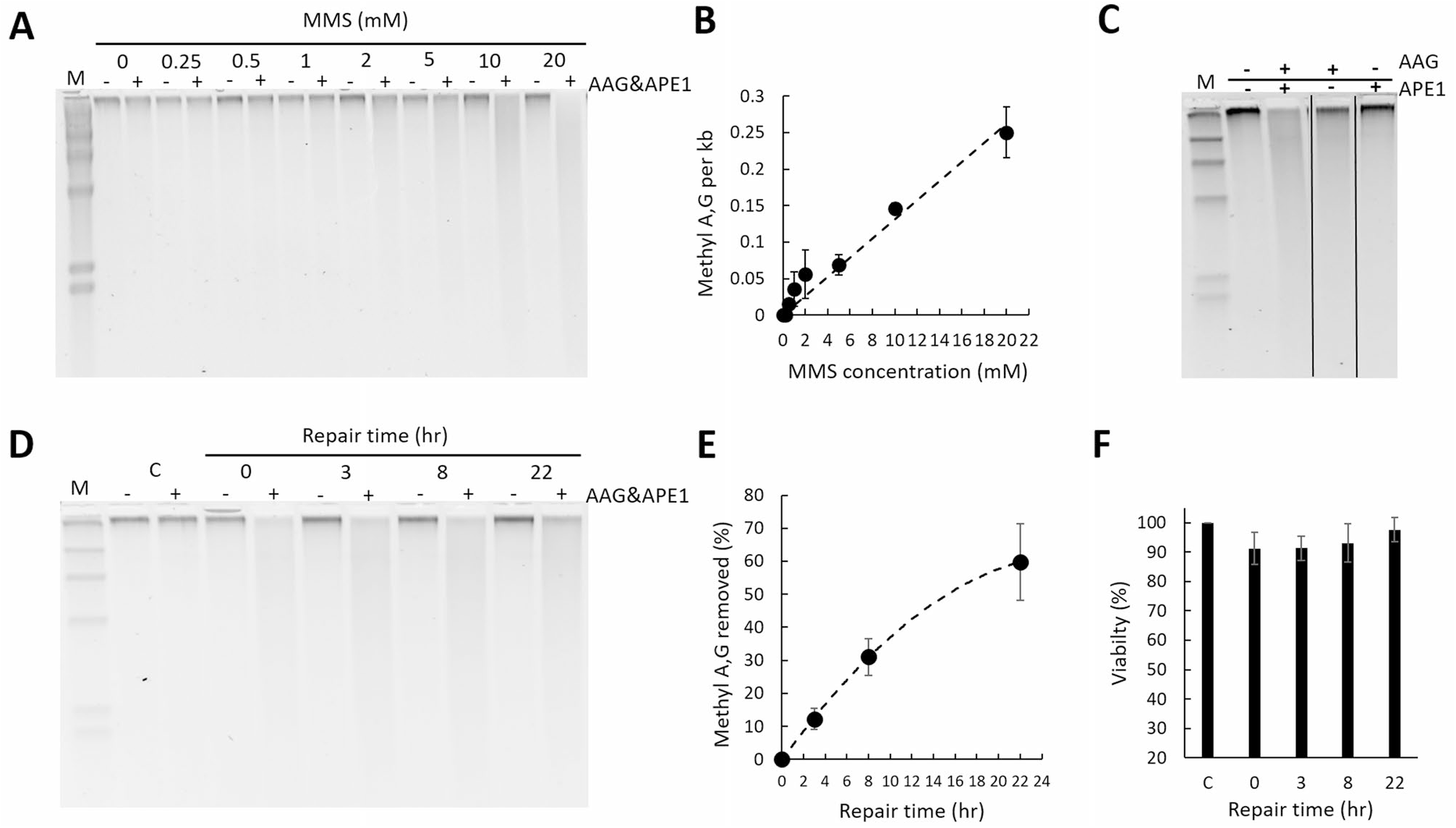
Alk-BER assay in human cells. (**A**) MMS dose response in SW13 cells. Cells were treated with increasing doses of MMS for 10 min at RT. Representative alkaline agarose gel image is shown. (**B**) Quantification of methyl A, G per 1 kb DNA fragment as a function of increasing MMS dose. The graph represents quantification of the data in panel (**A**). (**C**) Efficiency of double enzyme (AAG&APE1), and single enzymes: (AAG only), and (APE1 only), in converting methyl DNA adducts to SSBs. (**D**) Alkaline gel image representing DNA damage and repair time course. SW13 cells were exposed to 10 mM MMS for 10 min. Genomic DNA was isolated, and processed with double enzyme AAG&APE1 digest. (**E**) Quantification of the repair and removal of methyl A,G as a function of time. (**F**) SW13 cell viability measured by trypan blue. Each data point represents an average of 3 independent experiments; error bars were calculated based on standard deviation. Gel image presented in panel (**C**) has been cropped. Original, uncropped gel image is included in the supplementary data.

**Figure 5.**
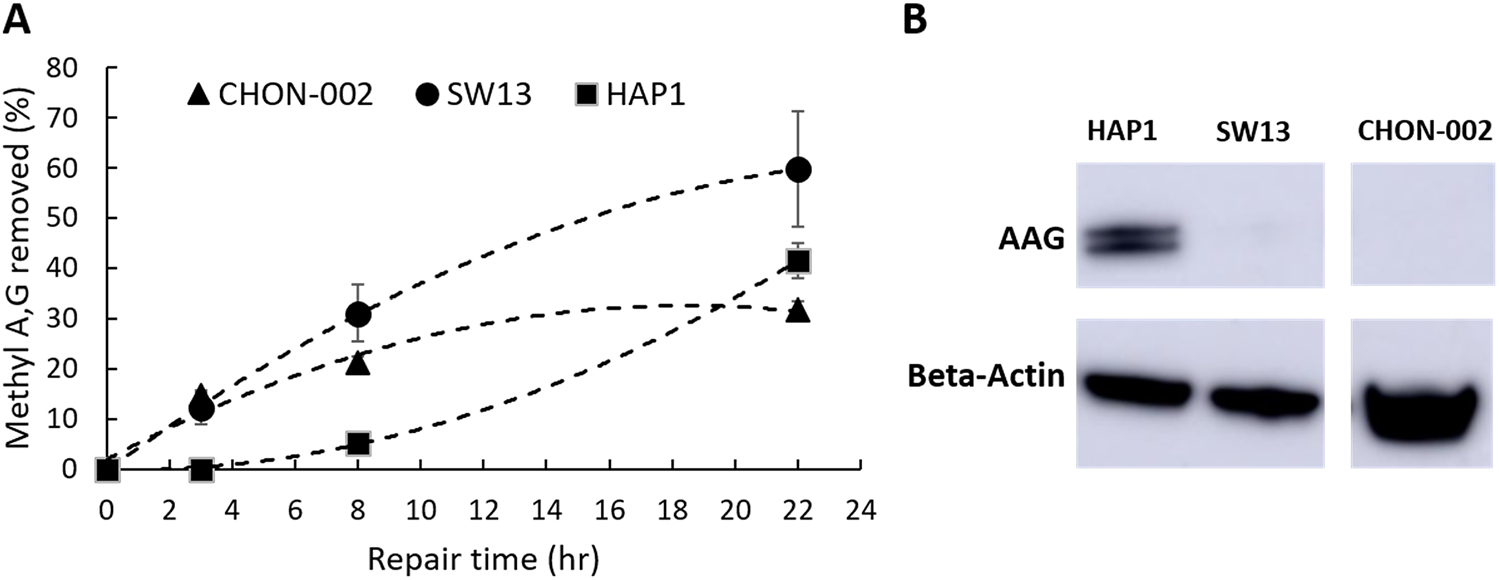
The repair of MDAs is slow in human cells and does not correlate well with the levels of endogenous AAG enzyme. DNA damage and repair time course experiment was performed in several human cell lines; CHON-002, SW13, and HAP1. Cells (60–70% confluent) were exposed to 10 mM (0.1%) MMS in 1xPBS for 10 min at RT, followed by DNA repair time course 0, 3, 8 and 22 h at 37 °C. (**A**) DNA repair rates were quantitated individually for each cell line and expressed as a percentage (%) of the removed methyl A,G, as compared to the 0 h time point. Each data point represents an average of two independent experiments. (**B**) Endogenous levels of AAG enzyme were detected by Western blotting. Western blot image presented in panel (**B**) has been cropped. Original, uncropped blot image is included in the supplementary data.

## Discussion

The efficiency of BER in repairing alkylation DNA damage varies substantially between different cells, tissues, and individuals, and has important implications in cancer development and treatment^1,2,58,59^. BER efficiency is a result of a complex interplay between genetic and epigenetic factors influencing the abundance and activity of the BER enzymes, and the individual steps in the BER pathway. Ability to measure the formation and rates of repair of alkyl DNA adducts in genomic DNA provides a direct assessment of BER efficiency in a given cell type.

Here, we report a quantitative cell-based assay, alk-BER (alkylation Base Excision Repair), developed for measuring efficiency and rates of BER following alkylation DNA damage in fungal model organisms (*S. cerevisiae, N. crassa*) and human cell lines (SW13, CHON-002, HAP1 and GM12878). Alk-BER offers a simple, time- and cost-efficient, cell-based method for quantitative analysis of alkylation DNA damage and repair in the genomic DNA. The alk-BER assay can be used to determine BER efficiency in various cell types, by assessment of the rates of methyl DNA adducts removal from the DNA over time.

Alk-BER has been developed as an adaptation and extension of the previous methods pioneered by Sutherland et al., to quantify UV-induced DNA damage and repair^48^. The original methods were based on quantification of DNA migration in ethidium bromide-stained alkaline agarose gels and scanning the image on film negatives with a densitometer. Alk-BER method reported here relies on the similar principles of quantifying DNA damage from alkaline agarose gels, but alk-BER assay has been developed and optimized specifically for the analysis of DNA alkylation repair. In addition, alk-BER involves several technological advances that improve method sensitivity and overcome many of the limitations of quantifying DNA from ethidium bromide fluorescence, including the linear range of detection. For example, we use SYBR Gold fluorescent dye to stain DNA in the gel. SYBR gold is about tenfold more sensitive than ethidium bromide^60^. SYBR gold also has increased signal to background ratio, as demonstrated by more than 1000-fold signal enhancement when bound to nucleic acids, which is approximately 50 times greater than that of ethidium bromide^60^. Furthermore, in our method gel images are acquired with modern, highly sensitive imaging systems (e.g., ChemiDoc MP, Bio-Rad) that allow for significant improvements of the linear dynamic range of detection. The improved linear range of detection permits accurate quantification of the DNA from a gel, where there is large variability in DNA band intensity (e.g. low intensity when bands disappear right after damage, and high intensity when bands reappear during repair)^51^. In summary, we adapted the principles of quantifying DNA damage from alkaline gel electrophoresis to analyze repair of DNA alkylation, and extended the method to analyze repair by BER at the genome level in various model organisms, including fungal cells (*S. cerevisiae, N. crassa*) and human cancer cell lines.

Yeast cells have been extensively used to study DNA repair processes in eukaryotic cells^61^. Yeast have a robust and conserved BER pathway to repair alkylation DNA damage. Using alk-BER assay, we found that in wild type strains of fungal model organisms (*S. cerevisiae* and *N. crassa*), BER proceeds quite rapidly, following removal of the MMS from the growth media. We showed that BER in the BY4741 strain of *S. cerevisiae* is nearly completed within 3–4 h post exposure since nearly 80–90% of the lesions were removed from the genome (Fig. 2C,D). Similarly, repair of alkylation DNA damage has been nearly completed within 2–4 h in the wild type laboratory strain of *N. crassa*^55^ (Fig. S2). We also showed that BER-deficient *mag1Δ* yeast mutant cells were deficient to clear MMS-induced lesions and accumulated MDAs over time, which validates specificity of the alk-BER assay (Fig. 3A–C). The human genome is much larger therefore repair is expected to take longer as compared to lower eukaryotes having smaller genomes. We found that the majority of the genome (∼ 70% lesions removed) was restored following 22 h post MMS exposure in SW13 cells (Figs. 4D,E, 5A), which is consistent with previous studies reporting that majority of DNA alkylation repair in mammalian cells can be completed within 24 h post exposure, as revealed by results generated with various quantification methods^39,62–66^. We found that the rates of BER to remove MMS-induced DNA adducts vary significantly between SW13 and CHON-002, and HAP1 cell lines and do not correlate well with the endogenous levels of AAG enzyme in these cell lines (Fig. 5A,B). These rates can be influenced by the endogenous levels and activities of various BER enzymes, including additional glycosylases, and other regulators of DNA repair. Future studies are needed to further investigate the mechanisms and regulators of DNA alkylation repair in human cells.

We found that in lower eukaryotes (*S. cerevisiae, N. crassa*), the rate of repair of MMS-induced methyl DNA adducts is strongly dependent on the functional MAG1 glycosylase, where *mag1Δ* mutants demonstrate abolished ability to repair MDAs. Interestingly, repair of MDAs in certain mammalian cells does not appear to be exclusively dependent on AAG glycosylase. It has been reported that the alkylated bases 3meA and 7meG, both AAG substrates generated from MMS treatment, are removed from the genome of AAG-deficient embryonic stem cells, with slower kinetics for 3meA but comparable kinetics for 7meG^67^. Other study revealed that similar levels of 7meG were detected in livers of AAG^+^/^+^ and AAG^−^/^−^ mice 24 h after exposure to MNU^68^. These studies suggest that in mammalian cells methyl DNA adducts can be excised and repaired in the absence of AAG enzyme, perhaps by involvement of other glycosylases, or spontaneous depurination. Future application of alk-BER could facilitate further understanding of the role of AAG and other factors in regulation of human BER.

The rates of repair of DNA alkylation can vary significantly between different organisms and cell types, and can be influenced by the levels and activities of the BER enzymes, genome size, chromosome landscape, DNA replication and translesion DNA synthesis (TSL). Therefore, it is necessary to experimentally determine a time range for accurate analysis of BER activity for each cell type under study. This is accomplished by carefully monitoring DNA damage induction and repair over a repair time course. The time range for BER to repair alkylation lesions is also dependent on the cell doubling time, which is much shorter (∼ 90 min) in yeast cells, and longer (∼ 16-24 h) in human cancer cell lines. It should be noted that exposure to DNA damage triggers transient cell cycle arrest as part of DNA damage response, that allows the time for DNA repair to take place before the next round of cell division^69^. The activity of BER should be analyzed immediately after induction of DNA damage, and within the 3–4 h for *S. cerevisiae and Neurospora*, and 24-48 h in human cell lines, when majority of the lesions are expected to be repaired by BER. After majority of lesions have been repaired, the DNA replication and cell cycle progression will resume. A small proportion of DNA base lesions, those which are left unrepaired, can be bypassed by translesion DNA synthesis (TSL) that occurs during DNA replication. TSL involves specialized replicative polymerases, which can perform error-free DNA synthesis over DNA base lesions^70^. Therefore, in order to avoid possible interference from DNA replication and lesion bypass, BER activity should be analyzed at early repair time points and within experimentally determined cell-specific time rage.

The alk-BER could serve as useful framework for number of approaches to study repair of DNA alkylation. For example, alk-BER assay could be used to distinguish, in a quantitative way, between the levels of MDAs and levels of downstream repair intermediates, such as AP sites. Highly specific and sensitive detection of AP sites could also be performed by processing of the sample with the AAG enzyme only (converts MDA to AP sites) and subsequent detection of AP sites using a highly sensitive AP site detection kit (e.g., Abcam, ab 211154). Alk-BER assay can also serve as a framework for quantification of gene-specific repair when coupled with Southern blot and hybridization of gene-specific probes. Alk-BER could also be useful in detection and quantification of MMS-induced methyl DNA adducts in preparation and optimization of samples for the approaches involving next generation sequencing, such as NMP-seq.

In summary, the alk-BER assay offers a versatile, reliable and affordable approach for quantitative analysis of DNA damage formation and repair following exposure to DNA methylating alkylating agents. The alk-BER assay can be easily optimized to be used in any type of cultured cells, and integrated with the existing approaches to study mechanisms regulating BER balance and capacity. The assay has the ability to detect imbalances in the activity of the BER process, that is highly relevant to cancer development and treatment. Quantitative analyses of DNA alkylation damage and repair using fungal genetic model organisms and human cell lines offer unique opportunities to identify novel, conserved regulators of BER.

## Acknowledgements

We thank Dr. Ann Bode for help with proof reading the manuscript. We would like to acknowledge Dr. John Wyrick for helpful discussions and intellectual contributions to this work. This work was supported by grant R01ES002614 from the National Institute of Environmental Health Sciences (NIEHS) (to M.J.S. and J.J.W.), R21ES028549 from NIEHS to W.C., and Faculty Research Grant from University of Georgia to W.C. The contents are solely the responsibility of the authors and do not necessarily represent the official views of the NIEHS, NIH.

## Author contributions

Conceptualization: W.C., P.M., M.S. Funding acquisition: W.C., M.S. Investigation: W.C., Y.L., P.M., E.Y.B. Validation: W.C., Y.L. Methodology: W.C., P.M., M.S. Project administration: W.C. Resources: W.C., M.S., Z.L. Supervision: W.C., M.S., Z.L. Visualization: W.C. Writing-Original Draft: W.C. Writing—review & editing: W.C., M.S., P.M.

## Additional information

### Supplementary Information

The online version contains supplementary material available at https://doi.org/10.1038/s41598-021-97523-w.

### Reprints and permissions information

is available at www.nature.com/reprints.

### Publisher’s note

Springer Nature remains neutral with regard to jurisdictional claims in published maps and institutional affiliations.

### Open Access

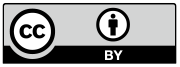

This article is licensed under a Creative Commons Attribution 4.0 International License, which permits use, sharing, adaptation, distribution and reproduction in any medium or format, as long as you give appropriate credit to the original author(s) and the source, provide a link to the Creative Commons licence, and indicate if changes were made. The images or other third party material in this article are included in the article’s Creative Commons licence, unless indicated otherwise in a credit line to the material. If material is not included in the article’s Creative Commons licence and your intended use is not permitted by statutory regulation or exceeds the permitted use, you will need to obtain permission directly from the copyright holder. To view a copy of this licence, visit http://creativecommons.org/licenses/by/4.0/.

